# High-content live-cell time-lapse imaging predicts cells about to die via apoptosis

**DOI:** 10.1101/2025.10.23.684203

**Authors:** Michael J. Lippincott, Jenna Tomkinson, Ibrahim Bilem, Mahomi Suzuki, Akiko Nakde, Toshiaki Endou, Simon Mathien, Felix Lavoie-Perusse, Carla Basualto-Alarcón, Gregory P. Way

**Affiliations:** Department of Biomedical Informatics, University of Colorado Anschutz, USA; Saguaro Biosciences, Quebec City, Quebec, CA; Yokogawa Electric Corporation, Tokyo, Japan; University of Montreal, Quebec, CA; Health Sciences Department, Universidad de Aysén, Coyhaique, Chile; Anatomy and Legal Medicine Department, Universidad de Chile, Santiago, Chile

## Abstract

Cell death is a dynamic process that unfolds through time. Live-cell time-lapse imaging captures these dynamics in a way that’s impossible for static snapshots. High-content imaging (HCI), which has been developed for static microscopy, applied to time-lapse imaging can quantify how single-cell states change through time. Here we show the ability of high-content live-cell time-lapse imaging (HCLTI) to quantify the onset and progression of one form of cell death called apoptosis. We apply the Live Cell Painting assay called ChromaLIVE^TM^ and develop an HCLTI analysis pipeline. We show that HCLTI can discern the morphology dynamics of cells undergoing apoptosis, and demonstrate that machine learning can predict apoptosis as early as 100 minutes after exposing HeLa cells to the apoptosis inducer Staurosporine. This technical advancement paves the way for future studies to better understand the dynamics of other forms of cell death. Understanding cell death dynamics is one piece of solving larger biomedical puzzles like understanding how cells resist death (e.g., therapeutic resistance of cancer cells) and how cells die too soon (e.g., neurodegeneration).

## Introduction

Regulated cell death (RCD), such as apoptosis, pyroptosis, ferroptosis, and necroptosis, are all complex cell processes that are crucial for the maintenance of complex life.^1–4^ When RCD is dysregulated (either upregulated or downregulated), disease arises, the processes of aging accelerate, and maintenance of life deteriorates. Understanding the properties, progression, and pathways of these RCD processes is critical for developing treatments for complex human disease. However, most knowledge about RCD is at the molecular level, for a few key molecules (e.g., caspase-1, caspase-3), and at single points in time. These approaches present challenges for understanding systems biology perspectives for complex biological processes.^5–8^

Additionally, single-molecule readouts do not capture cell states, and thus cannot capture potential pleiotropic effects. To better understand RCD, we must move beyond single-marker, static approaches and instead assay systems biology readouts through time.

We study RCD with a systems biology perspective through time to measure cell state in a scalable unbiased and minimally pleiotropic manner. We turn to the Live Cell Painting assay, ChromaLIVE^TM^, which is a non-toxic, multi-chromatic live-cell dye that passively integrates into lipid membranes, staining multiple cell structures and compartments.^9^ Due to its unique spectral properties and high sensitivity to intracellular environmental and physiological changes, the dye exhibits dynamic shifts in signal intensity and distribution, consistent with phenotypic changes.^9–11^ These phenotypic dynamics can be captured by HCI, specifically image-based profiling^12^ and quantified through time thereby enabling an unbiased, quantitative assessment of dynamic cell states. We refer to this approach as high-content, live-cell, time-lapse imaging (HCLTI) to more comprehensively chart complex biological processes.

Here, we applied ChromaLIVE^TM^ to profile over 20,000 single HeLa cells undergoing apoptosis in response to the apoptosis-inducing treatment Staurosporine at 10 different doses. We assayed these cells in 30-minute windows across six hours of imaging using the Yokogawa Cell Voyager CQ1 system. We developed an HCLTI analysis pipeline to extract and process CellProfiler computer vision features and scDINO deep learning features.^13,14^ We observed dramatic changes in cell morphology landscapes through time as cells undergo apoptosis.

Applying machine learning, we were able to predict cells that enter apoptosis in just 100 minutes after treatment with a pro-apoptotic agent. Our HCLTI work paves an important framework to expand the study of cell death dynamics (and all other dynamic biological processes) that unfold over time across different cell lines and disease contexts.

## Results

### Profiling apoptosis through time

We treated HeLa cells with ten different doses of staurosporine from 0 nM to 156.25 nM (**Figure 1A**). Staurosporine is a protein kinase inhibitor that invokes apoptosis via caspase-3 activation.^15–19^ We then applied the Live Cell Painting assay (ChromaLIVE^TM^)^9^, and performed live-cell time-lapse spinning disk confocal microscopy (Yokogawa CQ1), acquiring images per field of view (FOV) every 30 minutes in three z stacks for six hours. To establish the ground truth of which cells were undergoing apoptosis, we fixed the cells after six hours and performed immunohistochemistry (IHC) staining of AnnexinV to mark apoptotic cells^20^, and acquired images.

**Figure 1.**
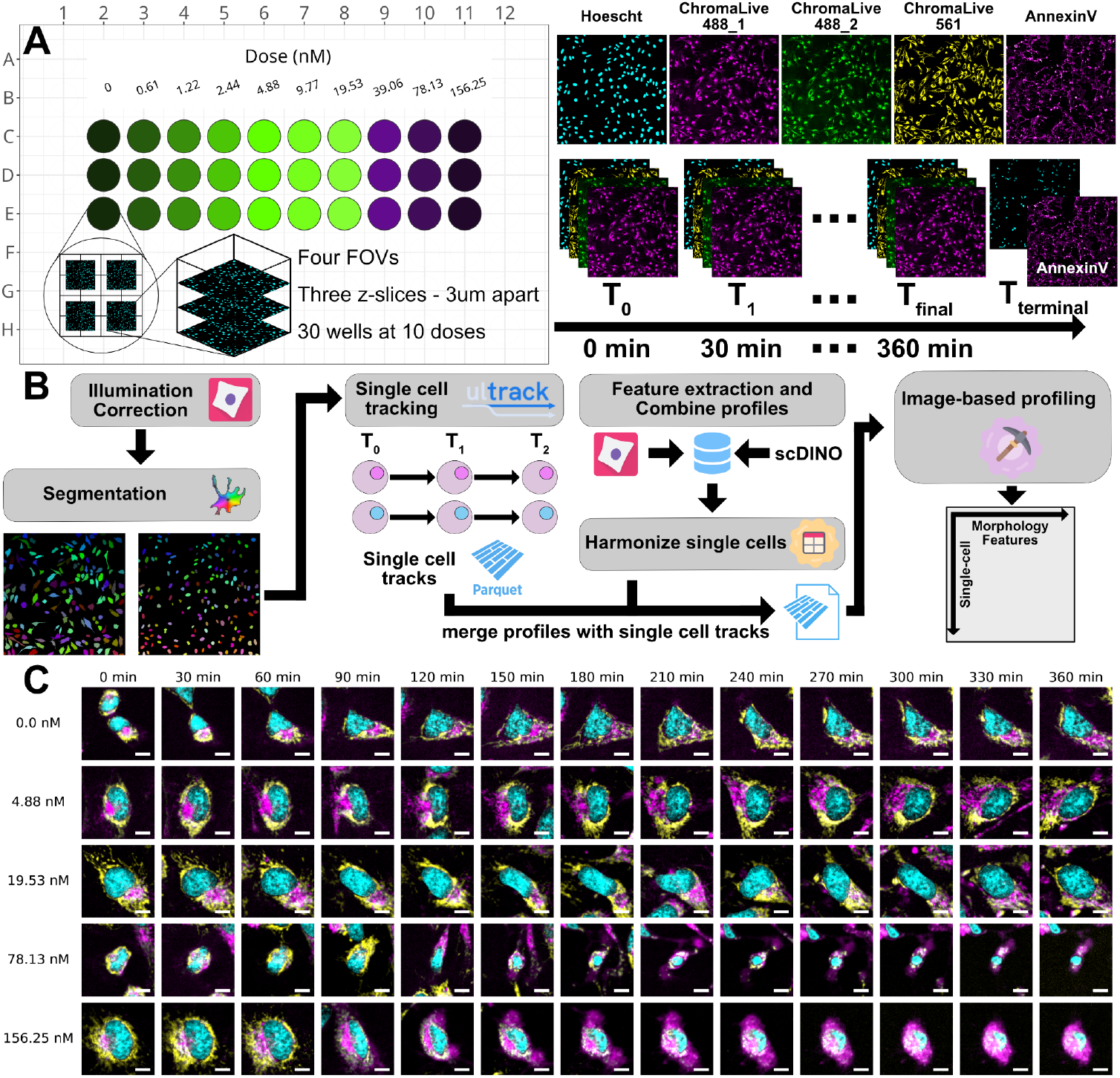
High-content, time-lapse imaging (HCLTI) workflow to profile apoptosis in HeLa cells. **(A)** The platemap showing replicates of ten different doses of staurosporine, an apoptosis-inducing compound. We imaged each well at four fields of view (FOVs) at three z slices, each 3um apart. Images were acquired every 30 minutes for 360 minutes (six hours), capturing Hoechst, ChromaLIVE 488_yellow, ChromaLIVE 488_red, and ChromaLIVE 561. We fixed cells at six hours and stained for AnnexinV, an apoptotic marker, and Hoechst. **(B)** Our HCLTI pipeline for processing time-lapse microscopy images. **(C)** Representative images of tracked HeLa single cells across time for varying doses of staurosporine. Cyan represents nuclei, yellow represents ChromaLive 561, and Magenta represents ChromaLive 488 at both its emission channels for visualization purposes only. Scale bars = 10 um.

### Developing a time analysis pipeline

We developed an HCLTI pipeline to process all images taken at each timepoint. The pipeline is similar to how we would process a single-timepoint HCI (e.g., Cell Painting^21^) experiment, with a few notable differences after cells are segmented (**Figure 1B**). Briefly, we maximally projected z-stacks, performed illumination correction, and segmented every single cell, nucleus, and cytoplasm using Cellpose and CellProfiler.^14,22^ We then performed feature extraction in two ways. First, we used CellProfiler to extract hand-engineered, computer vision features. Second, we used scDINO to extract classification (CLS) token embeddings of each single cell.^13^ These CLS token embeddings are a learned high-dimensional token in the DINO architecture, acting as a deep learning embedding.^23^ However, because our data are time-lapse, we diverged from the standard image-based profiling workflow.^10^ Specifically, we tracked single cells across time using the open-source Python package Ultrack.^24^ Ultrack assigns a unique ID to a single cell that is tracked through time, making it possible to relate profiled single cells across timepoints as paired data. After feature extraction, we merged single-cell profiles and single-cell tracks into a harmonized format using CytoTable^25^, and performed normalization and feature selection using Pycytominer.^26^ We normalized using the DMSO vehicle-treated cells at the first observed timepoint as a reference. In total, our time-lapse image-based profiles consisted of 188,065 total observations, with 20,677 single cells tracked through time (unique cells observed). Our two feature extraction pipelines measured 3,837 features (2,301 CellProfiler [CP] features and 1,536 scDINO embeddings). We performed feature selection on the concatenated (CP and scDINO) feature space, which resulted in 2,336 total features (800 CP and 1,536 scDINO). See the **Methods** section for complete data processing details. We subsequently generated aggregate profiles by calculating the median per feature of all single cells per well per time point.

Additionally, we generated consensus profiles, which are the median of all single cells across all replicate wells per time point, giving a profile representation for each dose of staurosporine.

### Profiling cells undergoing apoptosis reveals profound morphology changes through time

We applied our HCLTI pipeline to the apoptosis time-lapse dataset. We hypothesized that HCLTI would reveal how and when cells commit to cell death. To ensure that the main driver of phenotypic changes was unrelated to the cell count at each time point and dose, we imaged a relatively short drug exposure window of 6 hours. We confirmed cell count did not materially change across time with respect to all doses. Staurosporine did not yet yield substantial cell death, and the morphology we captured reflects cells actively undergoing apoptosis rather than dead and detached cells (**Figure 2A**). After applying our image analysis pipeline, we observed profound dose-dependent morphology differences across time and across multiple channels and feature groups (**Figure 2B; Supplemental Figure 2**).

**Figure 2.**
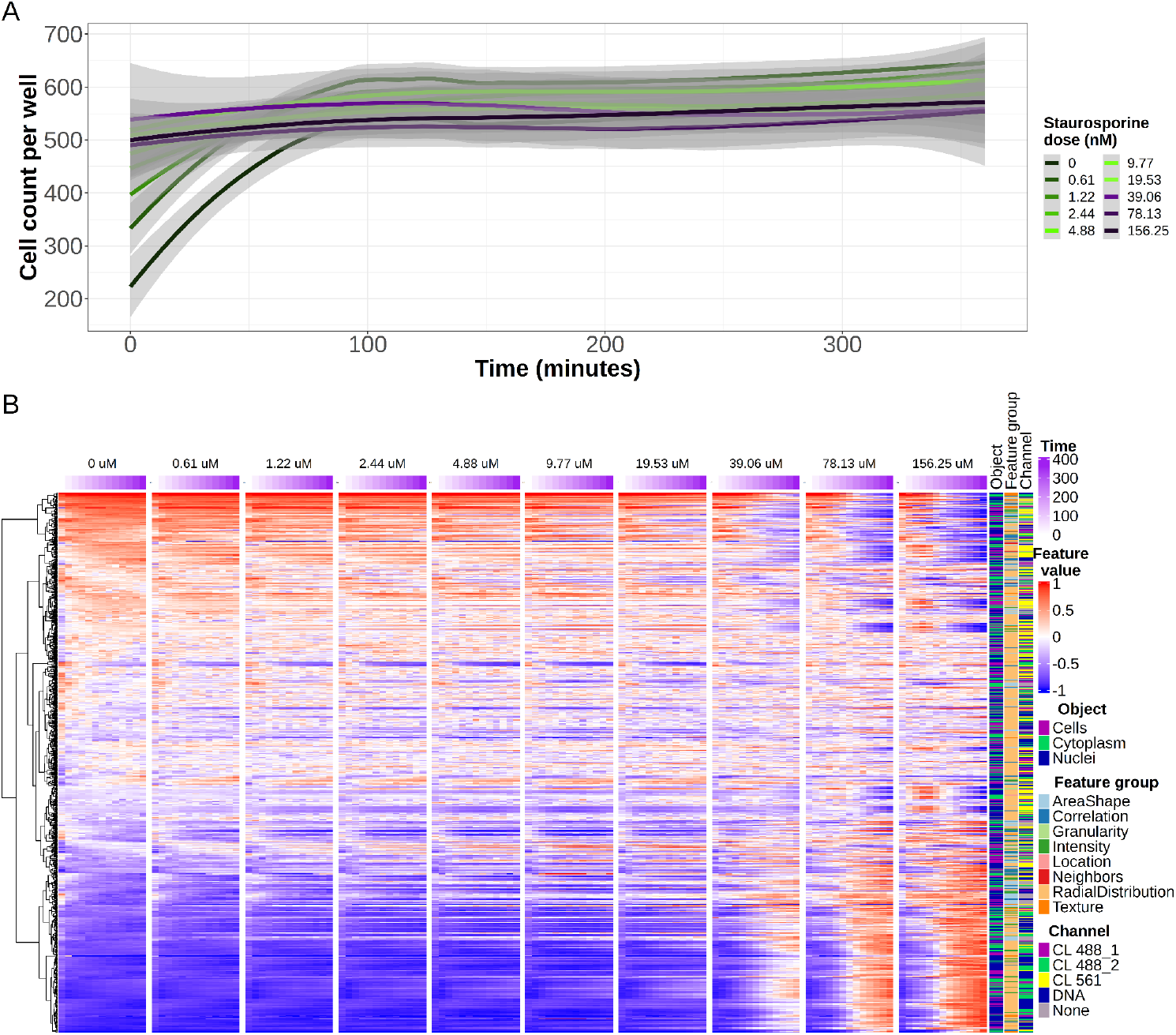
High-content, time-lapse imaging (HCLTI) of HeLa cells with increasing doses of staurosporine through time. **(A)** Counts of single cells within wells for each staurosporine dose and each time point. The grey boundaries per curve represent the standard error of cell count across replicate wells. **(B)** Consensus-level profiles of CellProfiler features across dose (blocks), and time (columns; colored on top by purple). We annotate each feature on the right by compartment (Nucleus, Cell, or Cytoplasm), feature group, and ChromaLIVE channel. We apply hierarchical clustering of CellProfiler features (row) based on only dose 0 nM of staurosporine (leftmost block). All other column blocks are ordered to this clustering solution.

We further investigated morphological differences through time by transforming single-cell profiles using Uniform Manifold Approximation and Projection (UMAP) (**Figure 3A; Supplemental Figure 3**).^27^ We fit the initial UMAP using all single cells of the lowest Staurosporine dose (0 nM - DMSO only), and transformed the remaining single cells into this space. We clearly see that single cells experiencing higher doses of staurosporine are substantially different than lower doses, which we summarize by displaying the average-density centroids per dose (**Figure 3B**). We next quantified the mean Average Precision (mAP) of mean aggregate profile replicates through time^28^, using timepoint 0 DMSO-treated well replicates as the reference control.^28^ We observed mAP near one for all treatments (except the lowest doses), which indicates high replicate reproducibility and consistently large morphological differences compared to the DMSO reference (**Figure 3C**).

**Figure 3.**
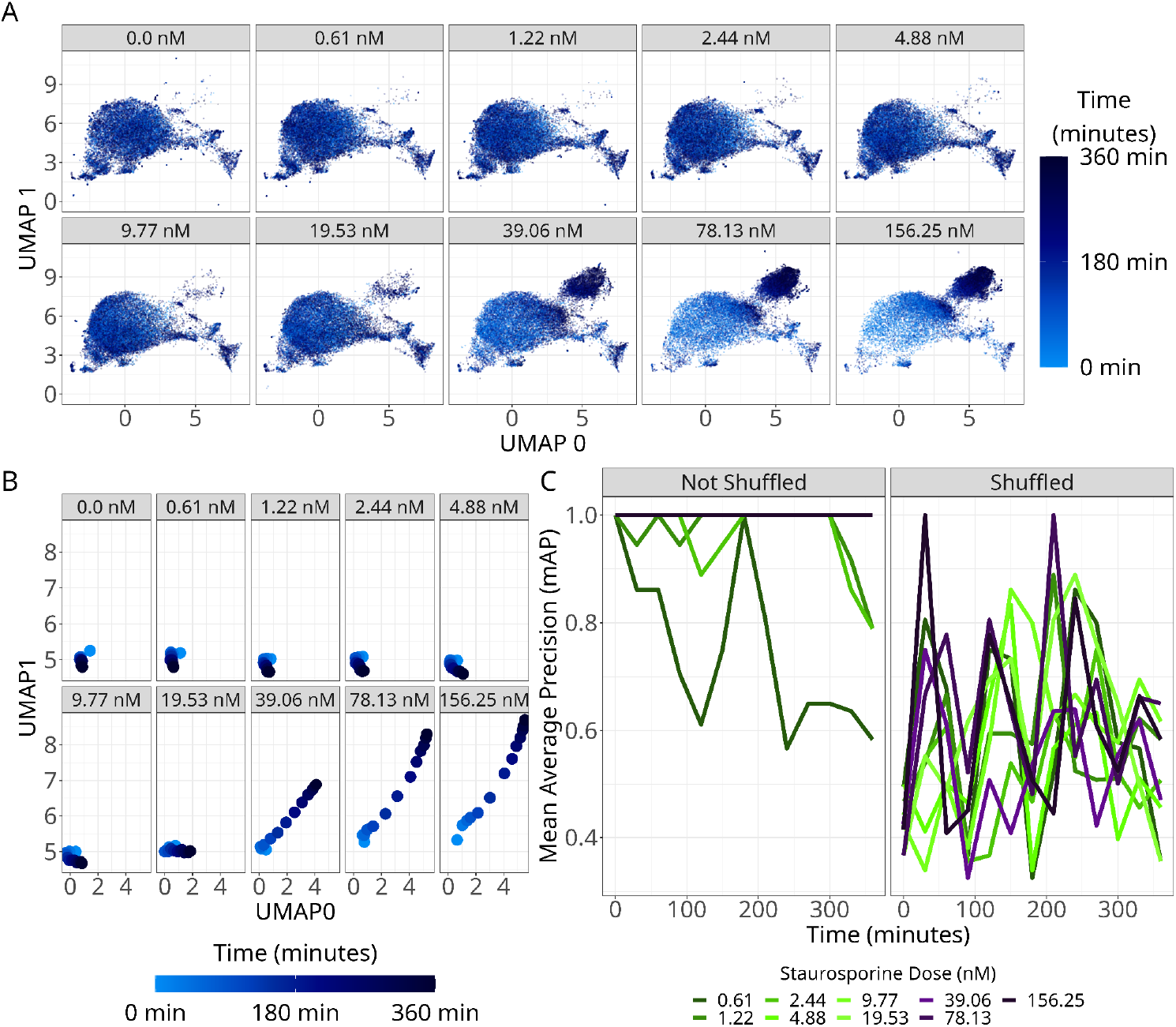
HeLa cells treated with an apoptotic agent change morphologies consistently and in dose-response through time. **(A)** Single-cell UMAP visualization through time (color bar) at ten different doses of staurosporine. **(B)** Projecting the average centroid of single cells per dose into UMAP space to summarize dose-specific changes through time. **(C)** Mean Average Precision (mAP) of aggregated profiles through time, per dose, using DMSO-treated wells at time zero as the reference. The shuffled panel indicates randomly permuted features prior to mAP calculation.

We next sought to quantify the influence of time, staurosporine dose, and cell density (confluence) on single-cell morphology feature variance. We fit 2,336 linear models (one per morphology feature), adjusting for the covariates of time, dose, and cell count per well. We found for ∼75% of features that time was the most important factor contributing to feature variance (**Figure 4**); however, these features shared significant variates with time as well with time only having 31 significant features and time plus dose having 402 significant shared features (**Supplementary Figure 4**). We see this trend for features across channels, cell compartments, and CellProfiler feature groups. We observe a similar trend for scDINO embeddings. We wanted to know the beta coefficients of covariate compared across featurization type, strictly as curiosity compelled us (CellProfiler vs scDINO). We found that 402 CellProfiler features significantly varied across time and dose, but only 51 scDINO features varied across time and dose (**Supplementary Figure 4B-C**).

**Figure 4.**
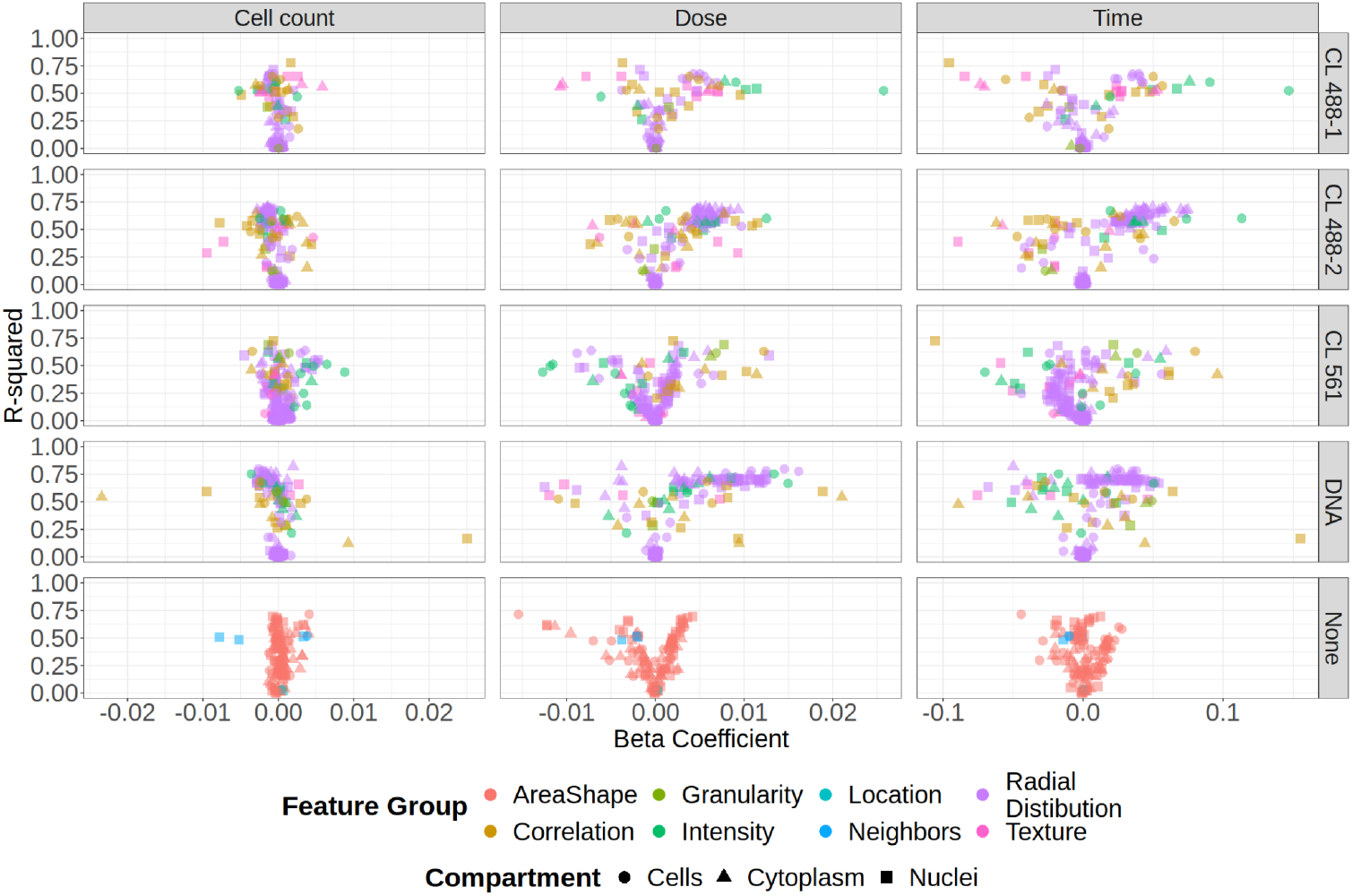
Linear modeling results adjusted by cell count, staurosporine dose, and time, separated by CellProfiler feature category. Linear model R-squared and beta coefficients plotted across channel, cell compartments, and CellProfiler feature groups.

### AnnexinV terminal staining is a ground truth apoptosis indicator in single cells

To confirm the presence of apoptotic cells, we fixed cells at six hours and stained them with AnnexinV to mark apoptotic cells and Hoechst to mark nuclear DNA (**Figure 5A**). As we expected, we observed an increase in whole-image median intensity of AnnexinV with increasing staurosporine dose (**Figure 5B; Supplemental Figure 5A**). We next performed an image-registration procedure to align the nuclear and cytoplasmic masks of the ChromaLIVE^TM^ final timepoint with the AnnexinV terminal timepoint. Using these masks, we applied a traditional image-based profiling pipeline to extract CellProfiler features from the annexinV and nuclear stains in nuclei, cells, and cytoplasmic compartments (**Supplemental Figure 5;** see **Methods** for details). Applying UMAP to the resulting single-cell profiles, we observed substantial single-cell heterogeneity and large differences across dose, with the most profound shifts starting at 39 nM (**Figure 5C; Supplemental Figure 5B**). In summary, AnnexinV confirms a dose-dependent morphological response to staurosporine with substantial single-cell phenotypic heterogeneity and low amounts of dead cells.

**Figure 5.**
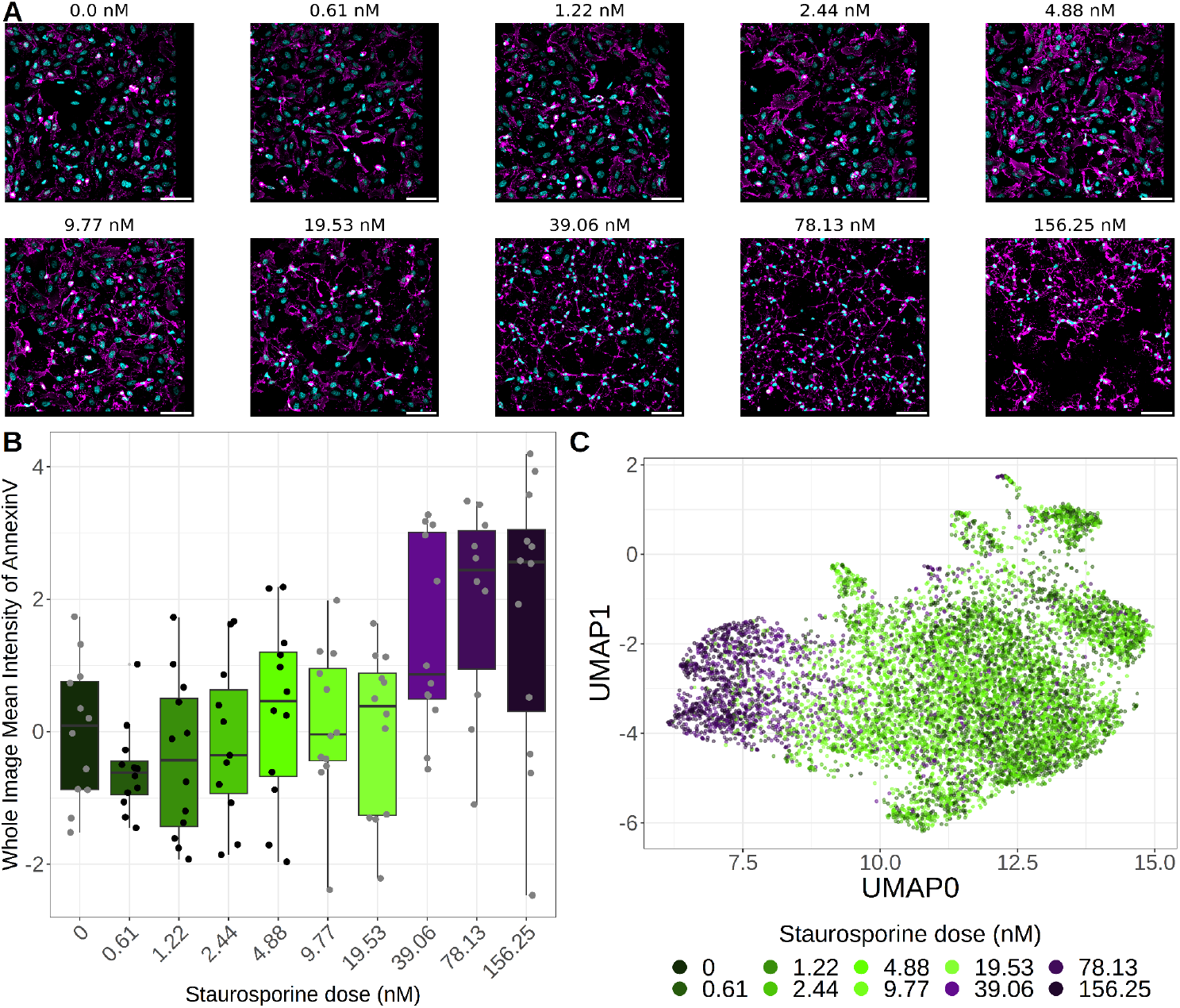
AnnexinV staining to establish single-cell ground truth of apoptosis. **(A)** Image montage of a randomly-selected field of view (FOV) from a single well of each of the 10 doses of staurosporine. Hoechst (cyan) marks nuclei, and AnnexinV (magenta) marks apoptotic cells. Scale bars represent 100 um. **(B)** Quantifying the mean AnnexinV intensity of whole FOV measurements across all wells. **(C)** UMAP of the single-cell profiles at the terminal time point (derived from AnnexinV and Hoechst features), colored by dose.

### Predicting the temporal dynamics of apoptosis

One major benefit of HCLTI is that it provides direct context of a cell’s past, present, and future. We hypothesized that we could predict at what point in time cells enter apoptosis or at least the point in time at which we can detect shifts in morphology that lead to higher amounts of AnnexinV. To test our hypothesis, we trained elastic net models to predict terminal AnnexinV. Specifically, using the aggregated profiles of the final ChromaLIVE^TM^ timepoint, we trained two models to predict: **1**) the full AnnexinV/Hoechst feature vector at the terminal time point, and **2**) only the AnnexinV integrated intensity within the cytoplasm. We also trained each model on a randomly shuffled (permuted) final timepoint as a negative control baseline. We observed high performance of these models, with mean squared error (MSE) less than 0.025 for both whole profile predictions (model 1) and predicting the annexinV integrated intensity within the cytoplasm (model 2) (**Supplemental Figure 6A**). Additionally, we observed R-squared values above 0.3 for whole profile prediction and above 0.6 for annexinV integrated intensity within the cytoplasm, indicating moderately-high performance (**Supplemental Figure 6B**). After training these models, we then applied each model to prior time points. Our aim was to determine if this approach could identify which time point a cell population can be detected as apoptotic (**Figure 6A**). We observe a clear, dose-dependent bifurcation of the prediction of the AnnexinV integrated intensity in the cytoplasm as early as 100 minutes, culminating in predictions at later time points that converge to ground truth (**Figure 6B**). Finally, we apply the whole profile prediction model (model 1) to profiles across time, and visualize predicted cell states in two principal components (56.83% of variance). We see that the higher doses converge to the actual terminal timepoint profiles as time progresses (**Figure 6C**).

**Figure 6.**
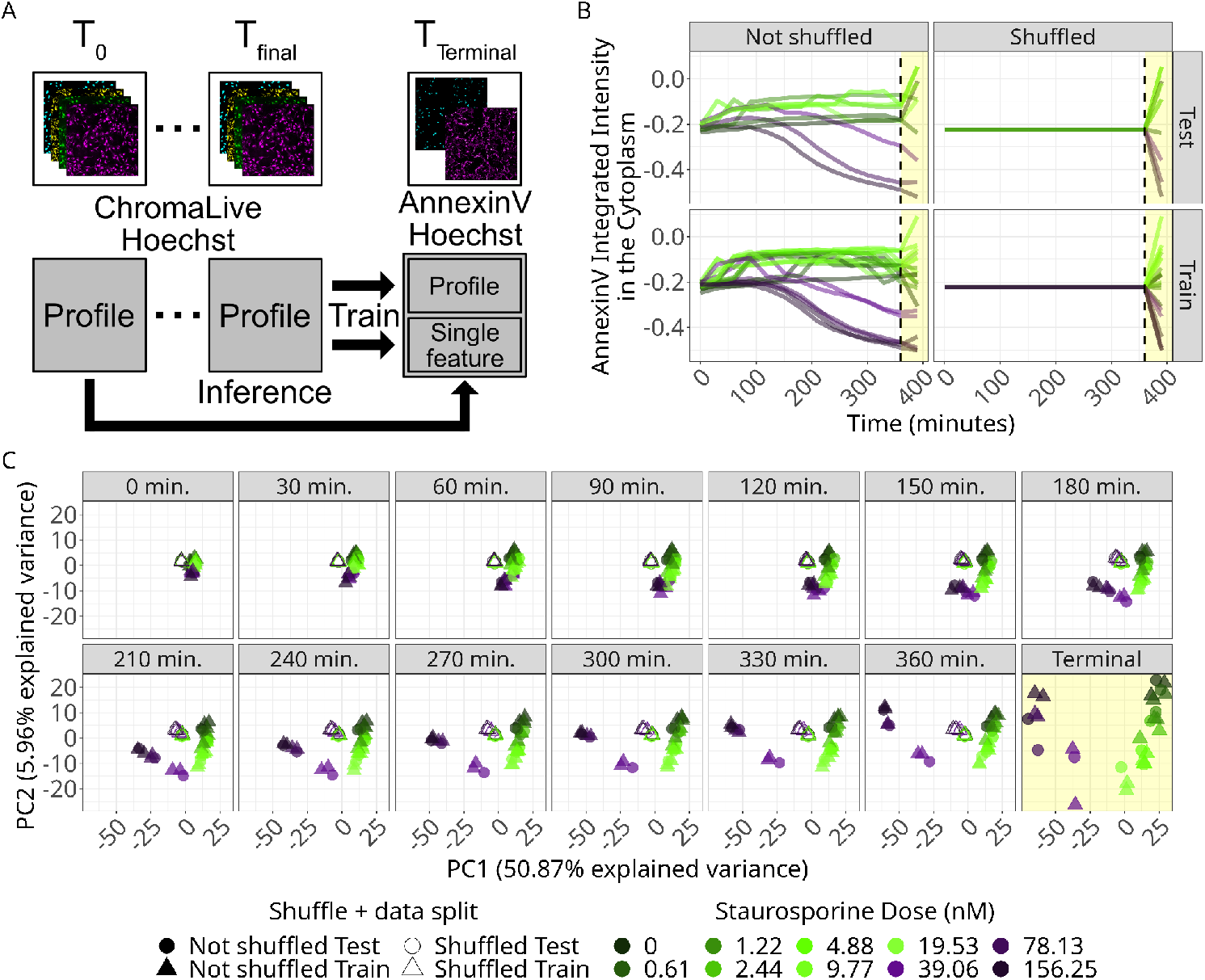
Predicting apoptosis through time. **(A)** Our machine learning workflow, where we trained on the final timepoint (containing ChromaLIVE^TM^ and Hoechst stains) to predict the terminal AnnexinV image-based profiles. We either trained a model to predict the full AnnexinV profile (model 1) or to predict the integrated intensity of AnnexinV in the cytoplasm (model 2) **(B)** Applying the trained AnnexinV terminal timepoint model to predict integrated intensity in the cytoplasm (model 2) at each prior timepoint. The ground truth annexinV integrated intensity in the terminal timepoint is highlighted in yellow. **(C)** The first two principal components of the predicted terminal image-based profile at each prior timepoint. Ground truth highlighted in yellow.

## Discussion

We treated HeLa cells with ten different doses of an apoptosis-inducing agent (staurosporine), applied the ChromaLIVE^TM^ assay, imaged with a live cell microscope, and developed a robust HCLTI pipeline to process single-cell, high-content morphology features through time. We also measured single-cell ground truth by fixing a terminal time point and marking apoptotic cells with AnnexinV. We identified profound and dose-dependent morphological differences as cells started to die via apoptosis. These differences were global, impacting all channels, compartments, and CellProfiler feature groups. These changes were mostly attributable to time and dose, rather than a marker of cell count-related differences. Importantly, we can predict an apoptotic cell population as early as 100 minutes after treatment, with each additional 30-minute interval improving prediction performance. In summary, we provide proof of concept of using HCLTI to reveal the morphological landscape of apoptotic cell death.

Previous studies quantify apoptosis via single-molecule assays such as the use of caspase cleavage and activation of apoptosomes.^1,29,30^ Using single-molecule readouts, while simple, can be pleiotropic, lack specificity, and reduce complex cell processes down to a single biochemical signal. In the modern digital age, advances in imaging and computational analysis now enable the generation of high-dimensional data that better reflect the complexity of biological responses. More recently, studies have used these quantitative and systems biology approaches to explore cell death processes^31,32^; however, most lack a temporal component and are conducted at a single timepoint. Some studies have used live-cell time-lapse approaches for apoptosis specifically^33–36^, but have lacked multiplexed morphological features provided by the Live Cell Painting and image-based profiling. Other studies have used image-based profiling approaches, but on fixed-cell assays, limiting their ability to capture time-sensitive phenotypic transitions.^36,37^ Other live-cell assays do exist,such as Acridine Orange staining, that provide valuable insights into the expression of a single molecular target. However, such approaches may not be fully compatible with prolonged live-cell imaging and do not capture the overall phenotypic landscape of cellular dynamics.^10^ All together, HCLTI proves to be a promising strategy for studying complex dynamic cell processes and for screening therapies that target these processes.

To study complex dynamic cell processes and adequately screen for new treatments for human disease, we need to meet the following requirements: **1**) the ability to observe a cell while it is alive, **2**) capture dynamic processes a cell carries out with respect to time, **3**) extract systems biology representations of a cell, and **4**) avoid survivorship bias. HCLTI can capture cells through time to sample its dynamic nature, and can have morphology measurements extracted. When studying cell death, the fourth requirement is arguably the most important. Depending on the compound and time point used in a static assay, the measurement or readout could be quantifying a cell’s reaction to a compound rather than the effect of the compound on the cell, which are distinctly different. To describe this difference, let’s start with a population of cells treated with theoretical compound X, a cytotoxic compound, for n hours. In this example, at 9 hours, we perform a static assay where most of the cells are dead. We are now measuring the compound’s effect on the cells (i.e., death), rather than the effect of how the cells respond to the compound, which may be profoundly different depending on the biological context (e.g., cell type). To decouple this nuance, we need a new perspective. We turn to HCLTI, where we can observe single cells through time undergoing dynamic cell processes and measure the effect of how the cells respond to a compound. We do not study biological problems related to spatial organization, such as nuclear localization of transcription factors, using 1D images (lines). Why then would we study spatial-temporal biological problems, such as cell death, with static 2D images?

Though HCLTI is robust, we observe multiple limitations in our work. First, we recognize that six hours of total imaging at 30-minute intervals is a relatively small time window; however, given the breadth of doses of staurosporine, we nevertheless observed a diverse number of cell states without cells dying. Additionally, we used only a single perturbation, staurosporine. This is one way to induce apoptosis and is in no way a complete representation of apoptosis induction. Other apoptosis induction modes target different pathways, have different temporal dynamics, and may intersect with other forms of cell death. For example, staurosporine broadly inhibits protein kinases, leading to both caspase-dependent and caspase-independent apoptosis mechanisms^16^. Thapsigargin affects calcium signalling and leads to the unfolded protein response, and subsequently apoptosis.^38,39^ Additionally, we only measured three replicate wells per treatment. This made our training and test data splits use single cells from only two and one well, respectively. Considering the size of the training split, we still observe relatively high performance of the models. Finally, we establish ground truth only at the terminal (six-hour) time point, and not throughout the live-cell imaging with an endogenous marker. A ground truth readout at the terminal time limits the ability to validate earlier time point predictions.

Future work will focus on cell death predictions of single cells through time instead of populations of single cells. Additionally, future work will expand to other forms of cell death with multiple mechanisms of induction for each across different cell lines. This future work will establish the Temporal Morphology Atlas of Cell Death, which will be a critical resource for quantifying the morphological cascades of different cells dying via different mechanisms. We can also expand HCLTI to study any biological process unfolding through time. Nevertheless, a key question that still remains: At what point in time does a cell commit to a cell death mechanism, and can we quantify the temporal heterogeneity of death commitment of every cell within a given population? Answering this question could unlock how to better sensitize specific cell populations for cell death commitment. This work paves the way to answer such questions by advancing multiple fields, including image-based profiling, cell death, live-cell time-lapse microscopy, and quantitative systems biology, by detecting one form of cell death, apoptosis, in real time from image-based profiles derived from Live Cell Painting.

## Methods

### Cell culture ChromaLIVE^TM^ staining

HeLa cells (ATCC® CCL-2™) were cultured in high-glucose Dulbecco’s Modified Eagle Medium (DMEM; Gibco) supplemented with 10% fetal bovine serum (FBS; Gibco) and maintained at 37 °C in a humidified incubator with 5% CO_2_. Cells were plated at a density of 1 x 10^4^ cells per well in a 96-well optical-bottom plate (Greiner Bio-One μClear®) in 100 μL of complete DMEM containing 0.1% (v/v) ChromaLIVE^TM^ dye (Saguaro Biosciences). The plate was incubated overnight under standard culture conditions.

Because ChromaLIVE^TM^ is non-toxic and non-fluorescent in the culture medium, becoming fluorescent only after entering cells, no washing steps are required prior to imaging, which facilitates its integration into automated, high-throughput workflows.

### Cell perturbation

On the following day, nuclei were stained, prior to Staurosporine treatment, by adding Hoechst 33342 (Thermo Fisher Scientific) to freshly prepared ChromaLIVE^TM^-containing medium at a final concentration of 0.5 µg/mL. After 15 min of incubation at 37 °C, Hela cells were treated with Staurosporine (Selleck Chemicals) for 6 hours in a 10-point dose-response series, prepared as 2-fold serial dilutions, with 156 nM as the highest assay concentration. Each condition was tested in triplicates and all treatments were delivered in 0.1% DMSO.

### Annexin V staining (ground truth)

Immediately after the 6-hour time-lapse, cells were stained with Annexin V-633 using the Annexin V Apoptosis Detection Kit (Nacalai Tesque, Japan). The culture medium was replaced with 50 μL per well of Annexin V Binding Buffer containing the Annexin V-633 reagent and the plate was incubated at room temperature for 15 minutes. Subsequently, 200 μL/well of binding buffer was added to each well prior to imaging.

### Image Acquisition

After treatment of staurosporine, we imaged FOVs on the Yokogawa CellVoyager CQ1 spinning disk confocal microscope with an Olympus UPLXAPO 20x dry lens with a 0.75 NA. Live-cell imaging of ChromaLIVE^TM^- and Hoechst-stained samples was performed every 30 minutes for six hours total. At the six hours endpoint, fixed and stained cells for AnnexinV and Hoechst were imaged. Four FOVs per well were imaged and each FOV included three z-slices spaced 3 um apart. Imaging settings are summarized in **Table 1**.

**Table 1.**
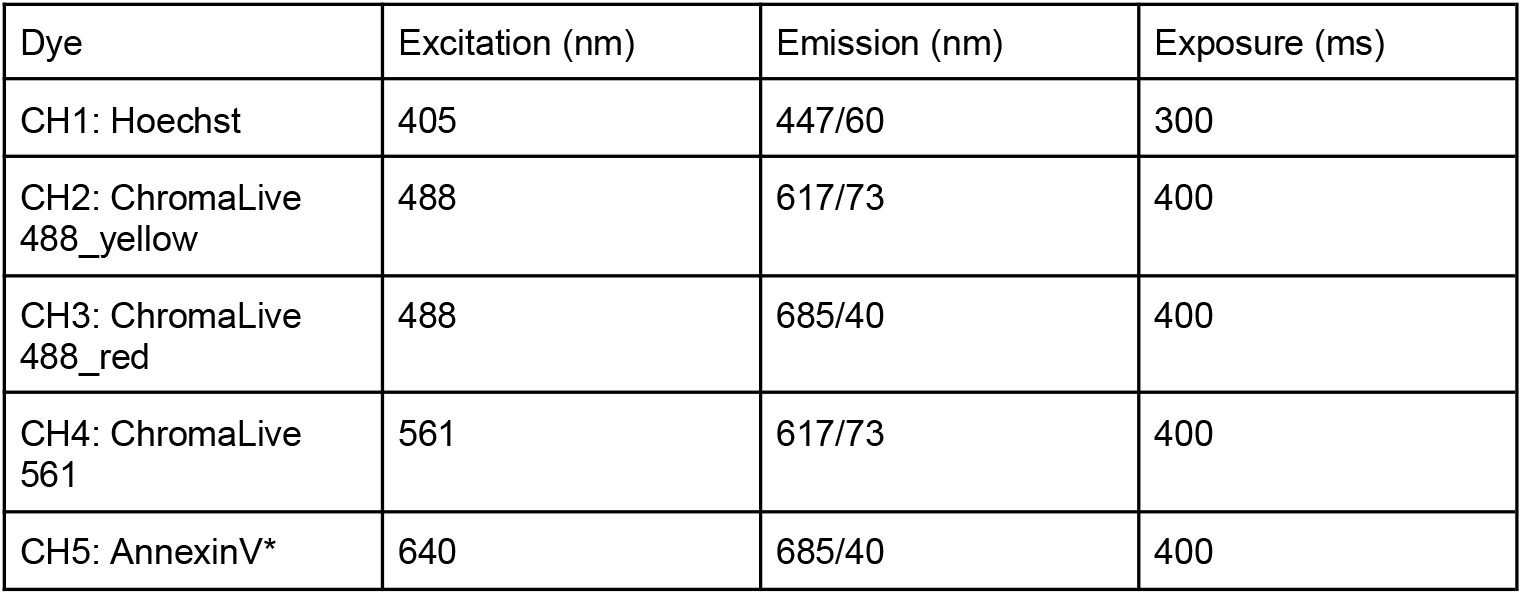
Imaging parameters used. *Note that AnnexinV was only used for a terminal staining and imaging step and not during the time-lapse acquisition.

| Dye | Excitation (nm) | Emission (nm) | Exposure (ms) |
| --- | --- | --- | --- |
| CH1: Hoechst | 405 | 447/60 | 300 |
| CH2: ChromaLive 488_yellow | 488 | 617/73 | 400 |
| CH3: ChromaLive 488_red | 488 | 685/40 | 400 |
| CH4: ChromaLive 561 | 561 | 617/73 | 400 |
| CH5: AnnexinV* | 640 | 685/40 | 400 |

### Image analysis: Illumination correction and segmentation

We performed a standard image analysis pipeline to process each individual frame in the time-lapse experiment. Specifically, we processed all images through an illumination correction pipeline using CellProfiler. We then performed nuclei segmentation using CellPose V, which utilizes the segment anything model.^22,23^ We performed cell and cytoplasm segmentation using CellProfiler with a propagation-based minimum cross-entropy segmentation algorithm.^14^ We saved masks for use in tasks such as image registration and feature extraction.

### Image analysis: time-lapse feature extraction

We used a standard CellProfiler^14^ feature extraction pipeline, with the exception of cell segmentation, to extract hand-drawn features for every single cell in all images across all time points. We used the CellProfiler profiles to find every single cell’s x,y centroid to generate single-cell crops (188,065). We then calculated the mean and standard deviation of the signal across every single-cell crop to generate a normalization tensor needed for scDINO.^13^ We then used snakemake to run the scDINO featurization pipeline. scDINO outputs a learned class (CLS) token that has 384 dimensions. Effectively, this gives 384 deep learning (DL) features per image. Given that we have four channels per single-cell crop, we extract (4 x 384) 1,536 deep learning scDINO embeddings. All workflows and scDINO pipelines were modified from the original scDINO GitHub repository from a fixed commit hash; links to GitHub repositories can be found in code availability.

### Image analysis: Image registration and alignment

We observed a slight x,y coordinate shift in the terminal images (containing nuclei and AnnexinV stains) compared to the final ChromaLive timepoint. To relate or track single cells across these time points, we performed image registration on the nuclei channels to register the offset of each FOV (**Supplemental Figure 7**). We then applied the specific offset to each of the terminal FOVs.

### Image analysis: Terminal segmentation and feature extraction

We imported the final ChromaLive timepoint cell segmentation masks to measure cytoplasm features in the terminal, AnnexinV images. We segmented terminal time point nuclei using the same CellProfiler-based approach we described for each ChromaLive time point. We used these cytoplasm and nuclei masks to featurize single-cells at the terminal time point. We also extracted whole image features from the registered and aligned terminal time point images.^14^

### Image analysis: Single-cell tracking

To track single cells across time, we used Ultrack^24^, a fast and scalable open-source scientific software for tracking cells in high-dimensional data. We output a single parquet file per FOV that contains multiple single-cell tracks. We merged these single-cell tracks with the generated profiles.

### Image-based profiling

We generated image-based profiles from every individual SQLite file output by CellProfiler and scDINO by using our open-source scientific software, CytoTable.^14,25,26^ We merged the single-cell tracks from Ultrack into the CellProfiler profiles prior to concatenating every single-cell CellProfiler and scDINO profile into one profile matrix (row = single cells, columns = metadata and morphology features). We next used our open-source software, Pycytominer^26^, to perform normalization of all single cells by fitting a standard scalar normalization (z-score) to the DMSO-treated cells at time point 0. The fit was then applied to all other wells and timepoints.

We subsequently used Pycytominer to perform feature selection using standard parameters. Specifically, we removed features that had: more than 5% of values that were NA or NULL; a Pearson correlation coefficient across all samples greater than 0.9; and low variance.

### Linear modeling

We performed linear modeling using the scikit-learn Python package.^40^ We fit one linear model per feature, adjusting for dose, cell count per well, and time. We recoded time to represent a linear step size of 1 between time points (e.g., time 0 min. = 1, time 30 min. = 2, etc.).

### Training elastic net models to predict AnnexinV

We trained two types of elastic net models. First, we trained an elastic net to predict the integrated intensity of AnnexinV in the cytoplasm using final timepoint profiles from the ChomaLive^TM^ and Hoechst profiles. Second, we trained an elastic net to predict the full terminal feature vector of AnnexinV and nuclear features using final timepoint profiles from the ChromaLive^TM^ and Hoechst profiles. We also trained negative control shuffled baseline models, in which we randomly permuted the final timepoint feature space (training data) to predict each endpoint. For every model trained regardless of input profile, we ran a five-fold cross-validation with hyperparameter optimization across the following hyperparameter space: alpha, [0.1, 1.0, 10.0, 100.0, 1000.0]; l1 ratio, [0.1, 0.25, 0.5, 0.75, 0.9]; with 10,000 max iterations. We trained each model using scikit-learn v1.6.1.^40^ We evaluated models only on the last timepoint to predict the terminal timepoint, we computed mean squared error and R-squared metrics on the testing dataset for both every model and its accompanying shuffled baseline model.

### UMAP and PCA Visualizations

We generated all single-cell Uniform Manifold Approximation Projection (UMAP^27^) plots using the following parameters: neighbors, 15; components, 2; distance metric, euclidean; random state, 42; minimum distance, 0.1; n_epochs, 500; learning rate, 1. We performed principal components analysis (PCA) by fitting on the whole dataset and visualized using the first two principal components.^41^

### mAP analysis

Mean Average Precision (mAP) is a rank-based metric that can be interpreted as a readout of how similar a given grouped replicate is to itself compared to all replicates within the reference control.^28^ mAP ranges from 0-1, where an mAP score of 1 would imply that a test group is more similar to itself than the control, and an mAP score of 0 would imply that a test group is virtually indistinguishable from the control group. We computed mAP using the copairs python package v0.5.0.^42^ We computed mAP for each time and dose combination using a reference from time 0 and dose 0 (the vehicle control). In addition, we calculated shuffled baseline mAP, where we randomly permuted the feature space at each distinct timepoint prior to mAP calculation.

### Computational resources

We utilized the Alpine high-performance computing resource at the University of Colorado Boulder. Alpine has 455 compute nodes with 28,080 cores, 36 NVIDIA a100 GPUs, 9 NVIDIA al40 GPUs and 36 AMD MI100 GPUs. In addition to Alpine, all other computation was performed on a local workstation with the following specifications: AMD Ryzen 9 5900x CPU—12 cores, with 128 GB of DDR4 RAM, and GeForce RTX 3090 TI GPU.

## Acknowledgments

This work utilized the Alpine high-performance computing resource at the University of Colorado Boulder. Alpine is jointly funded by the University of Colorado Boulder, the University of Colorado Anschutz, and Colorado State University, and with support from NSF grants OAC-2201538 and OAC-2322260.^43^ Data storage supported by the University of Colorado Boulder ‘PetaLibrary’.

## Data and Code Availability

Code for image processing, single-cell tracking, and image-based profiling can be found at: https://doi.org/10.5281/zenodo.17419422

Code for analysis, figures, and models can be found at: https://doi.org/10.5281/zenodo.17419412

We reference the scDINO repo: https://github.com/JacobHanimann/scDINO in which we modified files from a static commit hash: 3ad7adb05c64a1618c5ce3075a0ae756beff7800

## Glossary of Abbreviations

HCLTI: High Content Live-cell Time-lapse imaging
FOV: Field Of View
mAP: mean Average Precision
UMAP: Uniform Manifold Approximation Projection
CL: ChromaLive
CP: CellProfiler

