## Supplemental Figures for "High-content live-cell time-lapse imaging predicts cells about to die via apoptosis"

**Supplemental Figure 1. Representative images of tracked single cells across time and dose.** **2**
**Supplemental Figure 2. High-content scDINO features of HeLa cells change over time when treated with different doses of staurosporine.** **3**
**Supplemental Figure 3. Single-cell UMAP of each dose and timepoint of all single-cells comparing CellProfiler and scDINO feature spaces.** **4**
**Supplemental Figure 4. Linear model coefficients for scDINO features.** **5**
**Supplemental Figure 5. Image-based profiling workflow for extracting profiles in the fixed terminal time point.** **6**
**Supplemental Figure 6. Model performance of predicting terminal profiles** **7**
**Supplemental Figure 7. Registration pixel offsets per dose and field of view** **8**

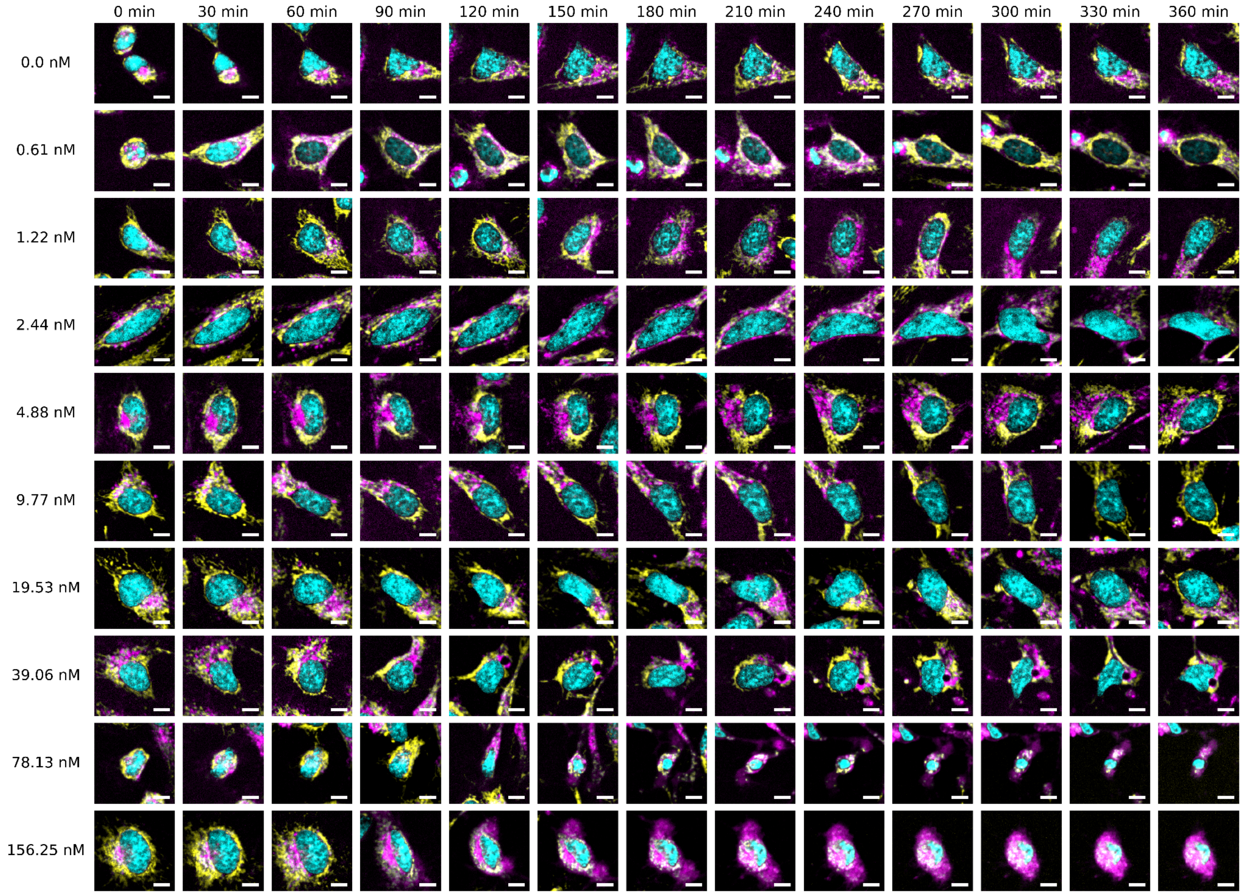


**Supplemental Figure 1.** Representative images of tracked single cells across time and dose.

We randomly selected a representative cell for each of the ten doses and tracked it over time. Cyan represents nuclei, yellow represents ChromaLive 561, and Magenta represents ChromaLive 488 at both emission spectra. Scale bars = 10 um.

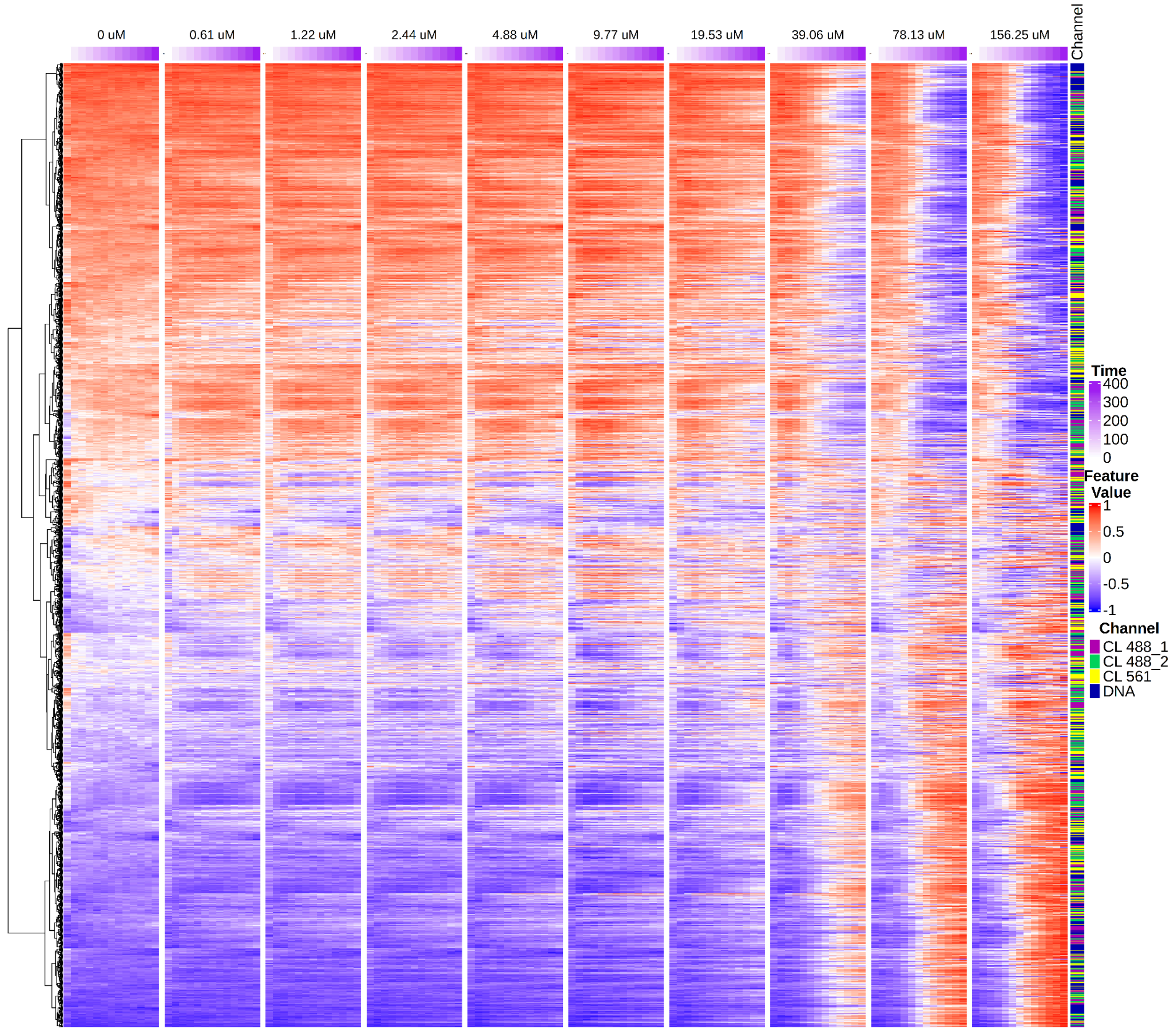


**Supplemental Figure 2.** High-content scDINO features of HeLa cells change over time when treated with different doses of staurosporine.

Related to Figure 2. Heatmap of both CellProfiler (CP) and scDINO profiles across staurosporine dose and time.

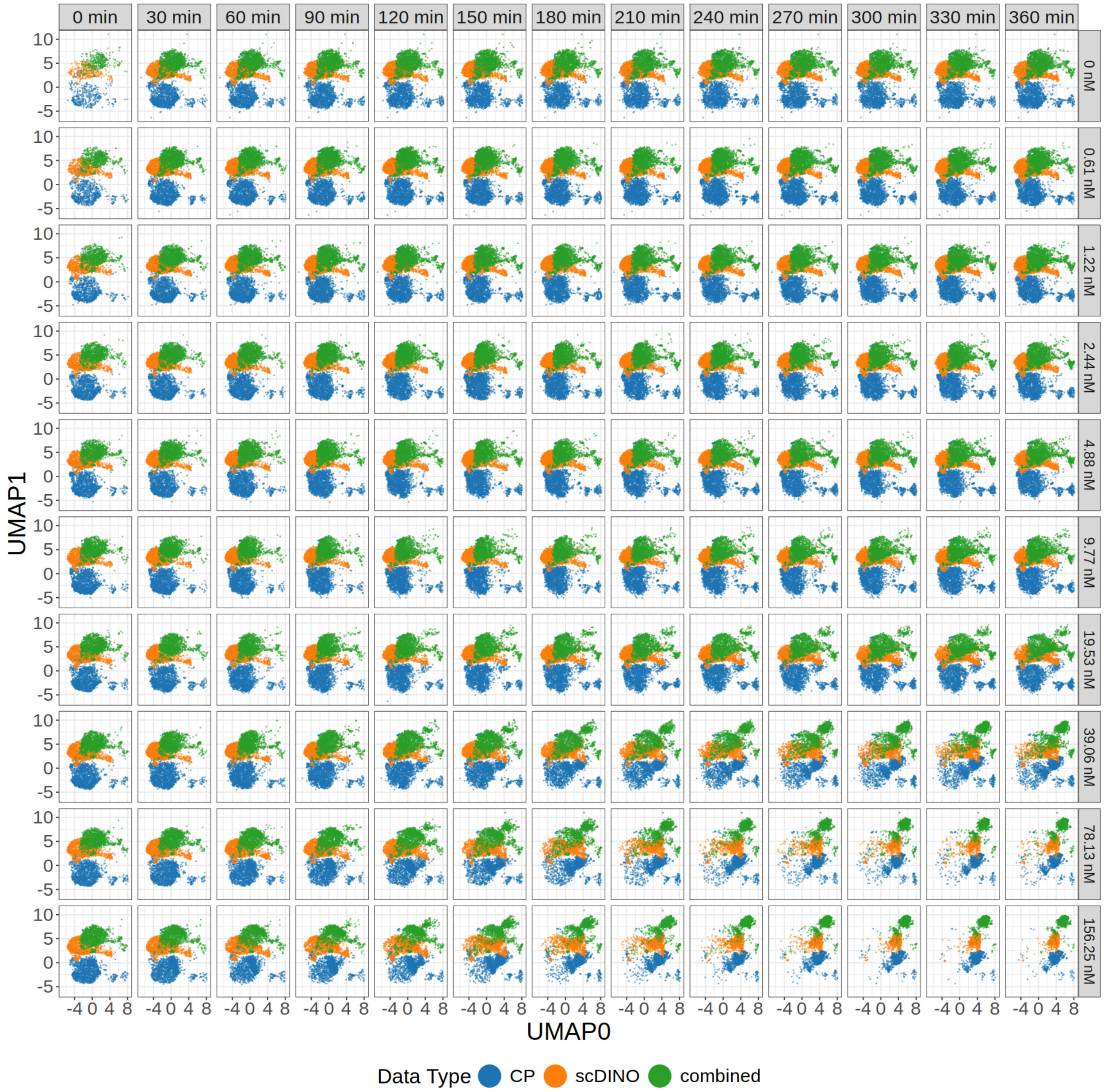


**Supplemental Figure 3.** Single-cell UMAP of each dose and timepoint of all single-cells comparing CellProfiler and scDINO feature spaces.

Time progresses from left to right and staurosporine dose increases from top to bottom. Each point is a single-cell. Each featurization method is indicated by color, CellProfiler (CP) in blue, scDINO in orange, and both feature spaces combined in green. UMAP originally fit on all cells from all timepoints.

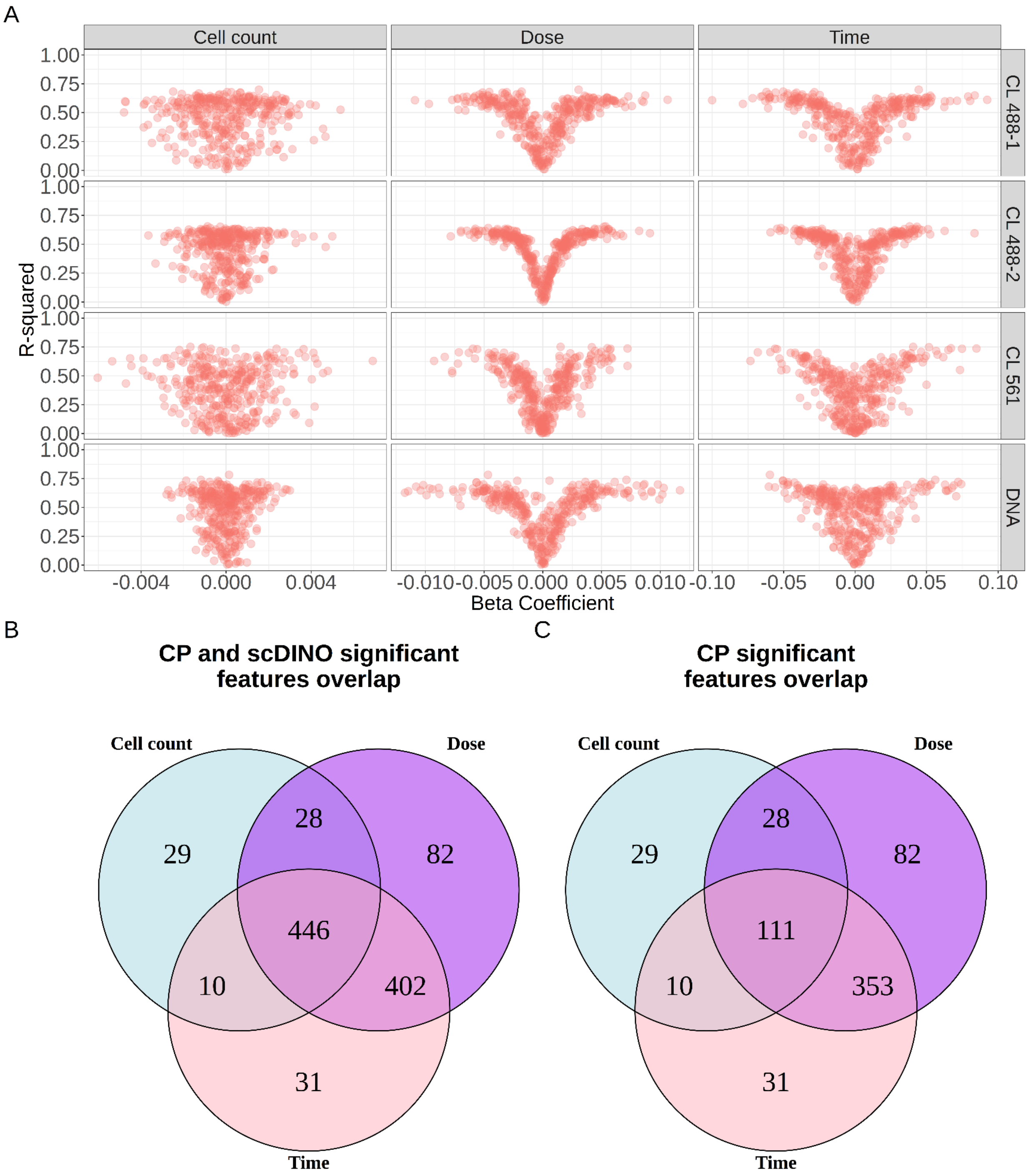


**Supplemental Figure 4.** Linear model coefficients for scDINO features.

**(A)** Beta coefficients for every variate in all feature models. Each variate is visualized by its feature model, separated by covariate, channel, and feature group. **(B)** scDINO and **(C)** CellProfiler (CP) features with significant contributions by cell count, dose, or time to feature variance.

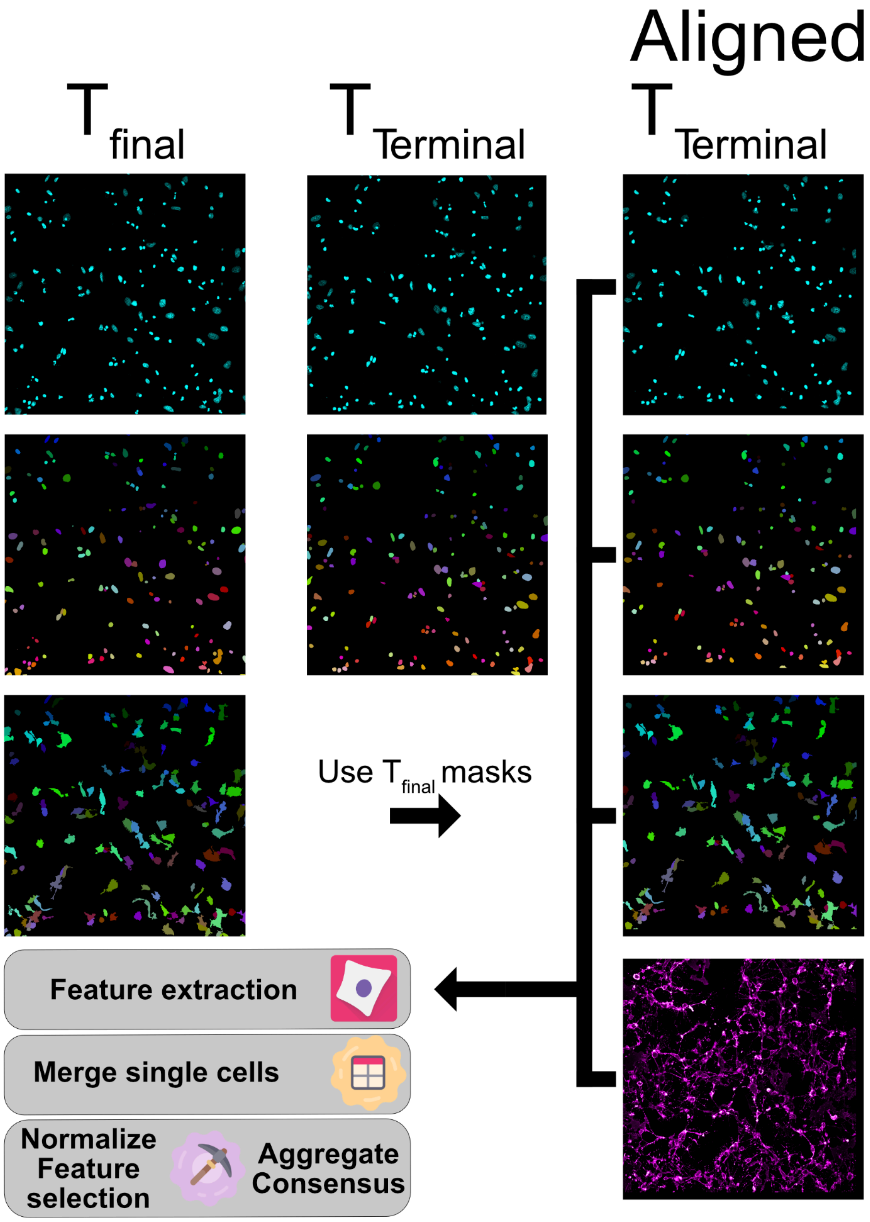


**Supplemental Figure 5. Image-based profiling workflow for extracting profiles in the fixed terminal time point.**

The fixed terminal time point does not include a cytoplasmic marker, so we could not readily apply cell segmentation. Instead, we performed image registration on the Hoescht channel between the final ChromaLive time point and  terminal timepoint. In effect, this generated an offset (x,y coordinate shift), which we applied to the rest of the terminal channels and masks for feature extraction.

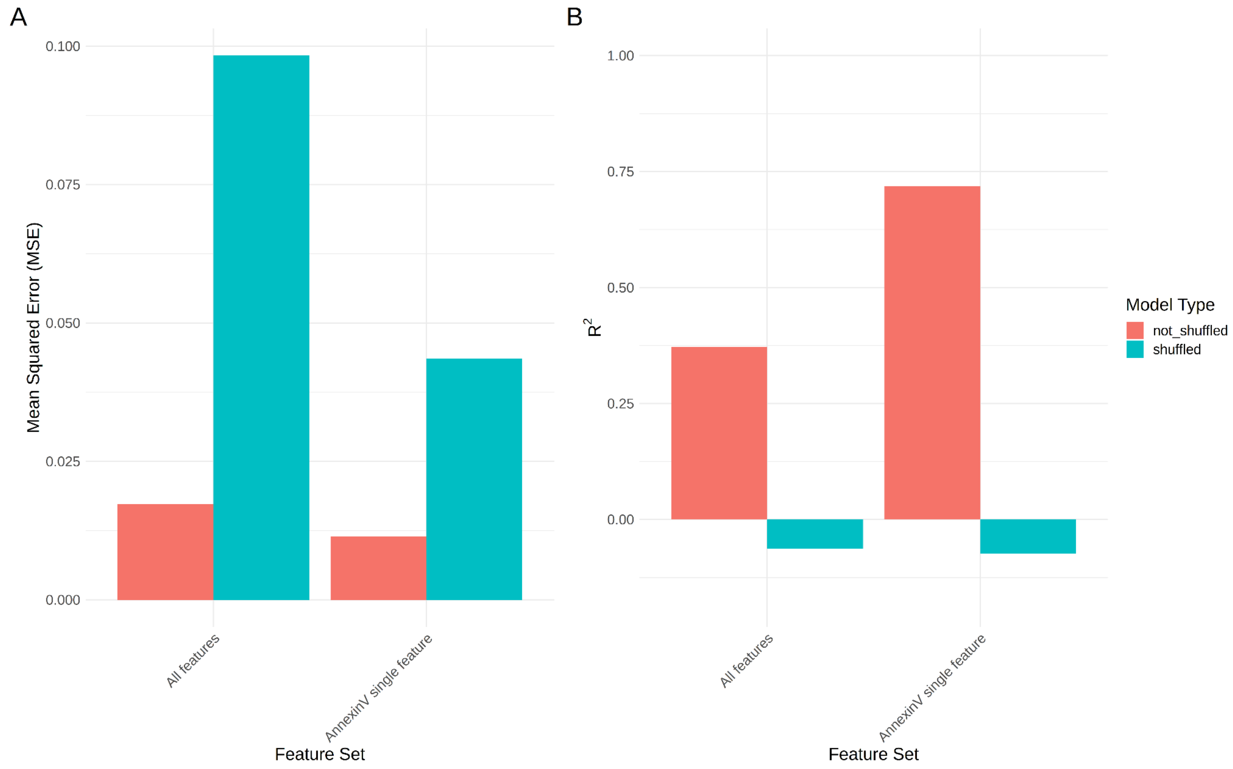


**Supplemental Figure 6.** Model performance of predicting terminal profiles

**(A)** The mean squared error (MSE) and **(B)** R-squared performance for predicting all terminal features and a single annexinV feature (Integrated Intensity in the cytoplasm) for both shuffled and non shuffled models.

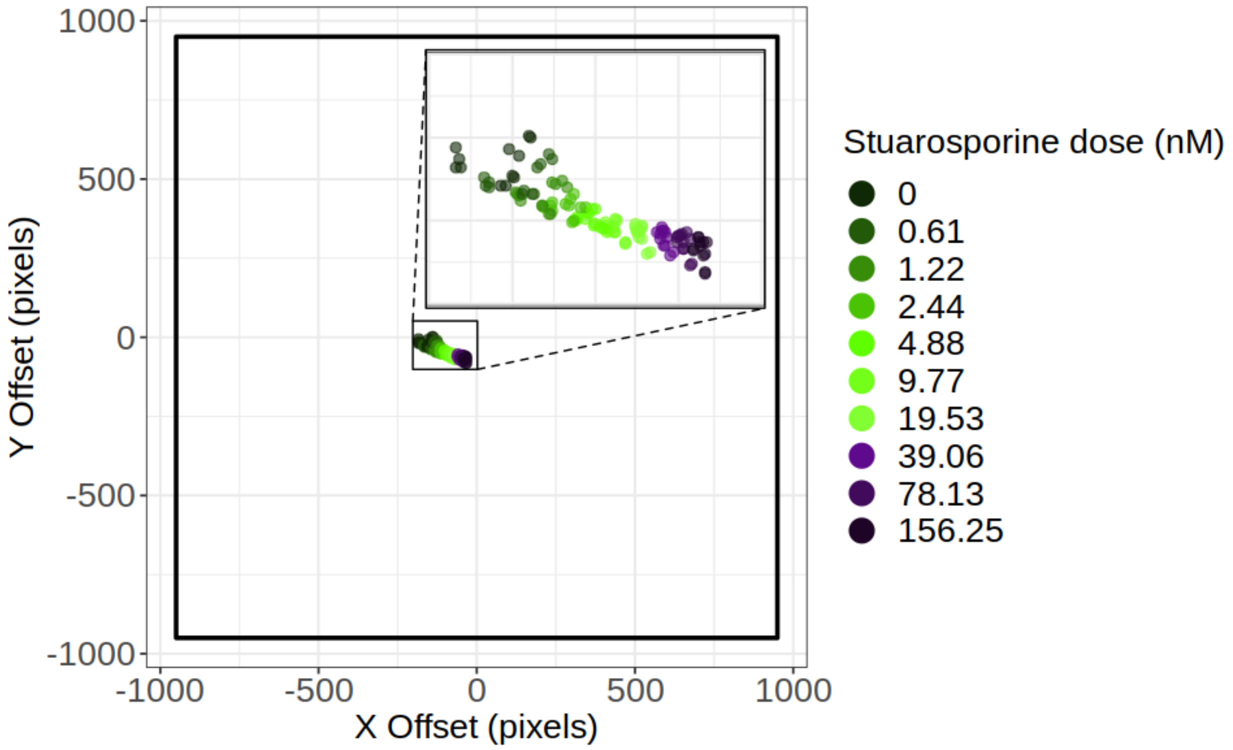


**Supplemental Figure 7.** Registration pixel offsets per dose and field of view

Cartesian coordinates of how far a single image (terminal time point) needs to be transformed to be aligned with a second paired image (ChromaLive final time point). Each image is 1900,1900 pixels; there are no offsets above 200 pixels. The black lines represent the image borders, and the offset is zoom to show consistency of plate layout relationship (doses sequentially increased toward right of the plate).
